# IFN-λ4 is associated with increased risk and earlier occurrence of gastrointestinal, respiratory and malarial infections in Malian children

**DOI:** 10.1101/2020.02.24.962688

**Authors:** Ludmila Prokunina-Olsson, Robert D. Morrison, Adeola Obajemu, Almahamoudou Mahamar, Sungduk Kim, Oumar Attaher, Oscar Florez-Vargas, Youssoufa Sidibe, Olusegun O. Onabajo, Amy A. Hutchinson, Michelle Manning, Jennifer Kwan, Nathan Brand, Alassane Dicko, Michal Fried, Paul S. Albert, Sam M. Mbulaiteye, Patrick E. Duffy

**Author notes:** Correspondence to: Ludmila Prokunina-Olsson, PhD, Laboratory of Translational Genomics, Division of Cancer Epidemiology and Genetics, National Cancer Institute, 9615 Medical Center Dr, Bethesda, MD 20892-9776.

## Abstract

Genetic polymorphisms within the *IFNL3*/*IFNL4* genomic region, which encodes type III interferons, have been strongly associated with impaired clearance of hepatitis C virus (HCV) infection. We hypothesized that type III interferons might be important for the immune response to other pathogens as well. In a cohort of 914 Malian children, we analyzed episodes of malaria, gastrointestinal and respiratory infections using information for 30,626 clinic visits from birth through up to 5 years of follow-up. Genetic polymorphisms *IFNL4*-rs368234815 and *IFNL3*-rs4803217 that functionally affect type III interferons were genotyped with TaqMan assays. For both genetic variants and each infection, we evaluated time-to-first episode and calculated odds ratios (ORs) for the risk of an infection episode during follow-up, controlling for relevant covariates. Compared to children with the rs368234815-TT/TT genotype (IFN-λ4-Null), each copy of the rs368234815-dG allele was associated with an earlier first episode of a gastrointestinal infection (p=0.003) and respiratory infection (p=0.045). The risk of experiencing an infection episode during the follow-up was also significantly increased with each copy of the rs368234815-dG allele – for gastrointestinal infections (OR=1.53, 95%CI (1.13-2.07), p=0.005) and malaria (OR=1.30, 95%CI (1.02-1.65), p=0.033). *IFNL4*-rs368234815 and *IFNL3*-rs4803217 were in moderate linkage disequilibrium in this population (r^2^=0.78), and all the associations for rs4803217 were weaker and lost significance after adjusting for rs368234815, implicating IFN-λ4 and not IFN-λ3 as the primary cause of these associations. We conclude that the ability to produce IFN-λ4 may have broad health-related implications by negatively affecting the immune response and clinical outcomes of several common infections.

## INTRODUCTION

The outcome of any infection is determined by a complex interplay between host and pathogen factors. One of the main host factors is the immune system, which can be modified by genetic polymorphisms in immune response genes. For example, variable ability to clear hepatitis C virus (HCV) infection spontaneously or after treatment has been significantly associated with polymorphisms in the genetic locus that encodes the family of type III interferons (IFNs) ^1–5^. Type III IFNs provide a localized immune response at the sites of pathogen entry, such as at the epithelial surfaces of the respiratory and gastrointestinal tracts, as well as in the liver ^6,7^. If this first line of defense provided by type III IFNs is successful, it decreases the necessity of a much stronger and systemic immune response provided by type I IFNs (IFNα and IFNβ) and type II IFN (IFN°C).

Humans have four type III IFNs, all encoded within the 70 kb region on chromosome 19. Three of these IFNs (IFN-λ1-3) can be inducibly generated in all the individuals, while IFN-λ4 is produced only in about 50% of the world population, including 90% of individuals of African ancestry, versus only 50% of Europeans and 10% of Asians ^5,8^. The ability to produce IFN-λ4 is an ancestral trait, which appears to be under negative selection in humans ^9^. The reasons for this selection are unclear but could be related to poor clearance of some deadly infections in the past. Thus, historic infections might have shaped the genetic landscape of this locus. On the other hand, the variable production of IFN-λ4 might have influenced and still be influencing susceptibility to infections and clinical outcomes.

The production of IFN-λ4 is controlled by a frameshift dinucleotide polymorphism, rs368234815-dG, within the first exon of the *IFNL4* gene. The open reading frame for IFN-λ4 protein is created by the dG allele, whereas the TT allele results in the truncated, non-functional protein (IFN-λ4-Null) ^5^. Another polymorphism, rs12979860, also known as the *IL28B* (*IFNL3*) marker, is located within the first intron of *IFNL4* and often used as a proxy for the functional rs368234815 due to population-dependent moderate to high linkage disequilibrium between these variants. Although rs368234815-dG is generally considered the causal variant due to its direct functional effect on the production of IFN-λ4 protein ^5^, additional polymorphisms in the *IFNL3/IFNL4* region may also demonstrate significant associations because of considerable linkage disequilibrium. One of these linked variants is a single nucleotide polymorphism (SNP) rs4803217, which is located within the 3’UTR of *IFNL3*; this variant was proposed to affect IFN-λ3 protein by modulating the stability of *IFNL3* mRNA ^10^.

The strong associations between the *IFNL3*/*IFNL4* genetic variants and HCV clearance have been well established and replicated by multiple studies. Additionally, a recent study reported an association between the *IFNL4* genetic polymorphisms (rs12979860 and rs368234815) and the clearance of respiratory RNA viruses in children from Rwanda ^11^. To further explore the hypothesis that type III IFNs might impact the immune response to diverse pathogens, we analyzed detailed health records of 914 Malian children and genotyped their DNA for the functional polymorphisms *IFNL4*-rs368234815 and *IFNL3*-rs4803217. For these children, information for 30,626 scheduled and walk-in clinic visits from birth through 1-5 years of follow-up was available for analysis.

We observed that compared to children genetically unable to produce IFN-λ4 (carriers of the rs368234815-TT/TT genotype, IFN-λ4-Null), those with rs368234815-dG allele were significantly more likely to have experienced at least one episode of gastrointestinal infection (p=0.005) and malaria (p=0.033) during up to 5 years of follow-up, with an earlier first episode of gastrointestinal infections (p=0.003) and respiratory infections (p=0.045).

The associations for *IFNL3*-rs4803217 were weaker and not significant after adjustment for *IFNL4*-rs368234815. These results suggest the role of type III IFNs, and IFN-λ4, specifically, in genetic control of the immune response to diverse pathogens of clinical significance.

## METHODS

### The Mali longitudinal birth cohort

We used cord blood samples and data collected by the longitudinal birth cohort study conducted in Ouelessebougou, Mali, between 2010 and 2016. The goal of the cohort was to explore host and parasite factors that influence susceptibility to malaria infection and disease during pregnancy and early childhood. The study enrolled pregnant women aged 15-45 years presenting for antenatal consultations at the Ouelessebougou Community Health Center (CESCOM). The eligibility criteria included a residency in the district of Ouelessebougou for at least one year before enrollment and absence of a chronic, debilitating illness. Malaria history or diagnosis was not an exclusion criterion. Newborn children of the participating women were enrolled at delivery and were followed-up for 1-5 years. Before enrollment, written informed consent was obtained from the parents/guardians on behalf of their children after receiving written and oral study explanations from a study clinician in their native language.

The protocol and study procedures were approved by the institutional review board of the National Institute of Allergy and Infectious Diseases (NIAID) at the US National Institutes of Health (ClinicalTrials.gov ID NCT01168271), and the Ethics Committee of the Faculty of Medicine, Pharmacy and Dentistry at the University of Bamako, Mali.

The study site was located 80 km south of Bamako, an area of intense but highly seasonal malaria transmission. The follow-up protocol included visits to the clinic scheduled monthly during the high malaria transmission season (June-December) and every two months during the dry season (January-May), as well as *ad hoc* walk-in clinic visits for any acute symptoms. Clinical information was collected during all visits by project clinicians and recorded using standardized forms.

Per study protocol, malaria infection was tested at each visit and was defined as positivity for *P. falciparum* malaria parasite on thick blood smear microscopy, with or without clinical symptoms. Clinical malaria was defined as fever >37.5°C, presence of malaria-related symptoms, and either a positive Rapid Diagnostic Test (RDT) for malaria antigens or visualized malaria parasites on thick blood smear microscopy. Severe malaria (SM) was defined as *P. falciparum* parasitemia and at least one of the following World Health Organization (WHO) criteria for SM - 2 or more convulsions in the past 24 hours, prostration (inability to sit unaided or in younger infants inability to move/feed), hemoglobin°C<°C5 g/dl, respiratory distress (hyperventilation with deep breathing, intercostal recessions and/or irregular breathing), or coma (Blantyre score°C<°C3). The presence of SM symptoms but with negative tests for malaria antigens or parasites was attributed to other diseases. All other parasitemia episodes, including those that were asymptomatic, were classified as non-severe malaria (NSM). For ever/never analyses, the NSM group included children with malaria history who never had an SM episode, while the SM group included children with at least one SM episode, regardless of the number of NSM episodes.

Gastrointestinal infections (GII) were defined by diarrhea; respiratory infections (RI) were defined by cough and chest-related signs and symptoms (sputum production, dyspnea, tachypnea, abnormal auscultatory findings). The etiology of these infections (bacterial, viral, parasitic, fungal) was not determined. All children received treatment as clinically indicated for their symptoms. Episodes of the same infection were considered independent if they occurred more than 28 days apart.

### DNA extraction and genotyping

Genomic DNA was extracted from cord blood clots using the QIAsymphony automation system with protocol Blood_200_V7_DSP and the DSP DNA Mini Kit (Qiagen) by the Cancer Genome Research Laboratory of the DCEG/NCI. The DNA samples were tested by microsatellite fingerprinting (AmpFLSTR Indentifiler, ThermoFisher) to confirm gender concordance with study records and exclude any mixed up or contaminated samples.

DNA samples were genotyped for *IFNL4*-rs368234815 and *IFNL3*-rs4803217 by custom TaqMan genotyping assays as previously described ^5,12^, (**Figure S1**). *HBB* polymorphisms rs334-T/A, *HBB*-Glu7Val (HbS, encoding hemoglobin variants A/A, A/S, and S/S) and rs33930165, *HBB*-Glu7Lys (HbC, encoding hemoglobin variants A/A, A/C, and C/C) were genotyped by Sanger sequencing as previously described ^13^, (**Figure S2**). Approximately 10% of all samples were genotyped in technical duplicates, which were 100% concordant for all markers. The distributions of rs368234815 and rs4803217 did not deviate from Hardy-Weinberg equilibrium (HWE). All genotyping and sequencing was done in the Laboratory of Translational Genomics (LTG, DCEG, NCI) blinded to demographic and clinical data. Of the 993 initial samples, 21 (2.1%) were excluded due to poor quantity/quality of the extracted DNA. Of the remaining 972 samples, 38 samples (3.9%) were excluded due to discordant *HBB* status between the study records (determined based on gel electrophoresis) and DNA testing, 18 samples (1.9%) were excluded due to sex discordance between study records and Identifiler profiles and 2 samples (0.2%) – due to failed rs368234815 genotyping. The final set included 914 children with available genotype, demographic and clinical data for 30,626 clinic visits; 447 (48.9%) and 467 (51.1%) of children were females and males, respectively. The children were followed-up from birth for an average of 147 weeks (range 52-264 weeks), with an average of 33.5 visits per child (range 5-81 visits).

### Statistical analysis

Haplotypes and linkage disequilibrium metrics (D’ and r^2^) between the markers were analyzed and plotted using Haploview 4.2 ^14^. The associations for *IFNL4*-rs368234815 and *IFNL3*-rs4803217 were tested based on additive and genotypic genetic models, using the TT/TT genotype or TT allele (IFN-λ4-Null) for rs368234815 and G/G genotype or G allele for rs4803217 as reference groups.

The occurrence of infection episodes (never/ever) was analyzed for both polymorphisms separately and in the same model using binary logistic regression models and adjusting for the follow-up time (weeks). *HBB* status coded as 0, 1 and 2 for AA (wild-type) vs. sickle-cell trait (A/S or A/C) vs. sickle-cell disease (S/S, S/C or C/C) was used as a covariate for all malaria-related analyses; sex and birth month (as a proxy for seasonal effects) were tested but not significantly associated with any outcomes (data not shown). Logistic regression models were used to evaluate the associations of *IFNL4*-rs368234815 with infections of interest adjusting for the occurrence of other infections (ever/never) to examine whether these associations were specific to particular infection types.

Kaplan-Meyer plots and Cox proportional hazards regression models were used to analyze time-to-first episode of each infection, with P-values calculated per rs368234815-dG and rs4803217-T alleles. The number of infection episodes during follow-up was analyzed using Poisson models with overdispersion (Quasi-likelihood), adjusting for the length of follow-up as an off-set in the model (offset = log(LastVisitAge)).

Since the study did not have permission to perform genome-wide analysis, we used self-reported paternal tribe as a proxy to control for possible population stratification. Of the 21 tribes, four tribes accounted for 92.5% of all the children: Bamana - 72.3%, Peulh/Fulani - 9.2%, Soninke - 6.0%, and Malinke - 5.0%. The reported paternal tribe (for all 21 tribes) was used as a random effect variable for logistic regression analyses (glmer package), Cox regression analyses with gamma frailty, and the analyses of infection episode counts per year (glmmPQL package). All the analyses were performed using specified packages in R (https://cran.r-project.org/). The results were not adjusted for multiple testing.

## RESULTS

### Analysis of infection episodes in the Malian children

We analyzed information for 914 children based on health records for each of the 30,626 routine and walk-in clinic visits from birth through up to 5 years of follow-up. During follow-up, 76.1% of children had a malaria episode, including 11.1% children that had at least one SM episode (regardless of NSM episodes) and 65.0% of children that had only an NSM episode (and never an SM episode); 88.1% and 97.7% children experienced a GII and RI episode, respectively.

Episodes of any infection (malaria, GII or RI) were reported at 12.7% of the routine and 76.3% of the walk-in visits (Table 1). Thus, most infection episodes were acute and sufficiently severe to require an emergency visit to the clinic without waiting for the next regular scheduled check-up visit. The overall frequency of malaria infection at any visit was 10.6% (5.4% at routine and 19.9% at walk-in visits); GII was reported at 8.4% of all visits, (1.8% at routine and 20.2% at walk-in visits), and RI was reported at 21.0% of all visits (6.5% at routine and 47.4% at walk-in visits). No infection was reported at most visits (n=19,808, 64.7%), while one and two infection types were reported at 9,407 (30.7%) and 1,380 (4.5%) visits, respectively. Only 31 visits (0.1%) reported all three infection types at the same visit.

**Table 1.** Distribution of clinic visits for 914 children from the Mali birth cohort study

|  | Clinic visits, n |  |  |  |
| --- | --- | --- | --- | --- |
|  | All visits# | Routine | Walk-in | Other |
|  | 30,626 | 18,654 | 10,884 | 1,088 |
| Visits reporting any infection, n, | 10,818 | 2,377 | 8,303 | 138 |
| % of all visits | 35.3 | 12.7 | 76.3 | 12.7 |
| Visits with malaria, n, | 3,260 | 1,015 | 2,167 | 78 |
| % of malaria visits | 100.0 | 31.1 | 66.5 | 2.4 |
| % of all infections | 10.6 | 5.4 | 19.9 | 7.2 |
| Visits with GII, n | 2,563 | 342 | 2204 | 17 |
| % of GII visits | 100.0 | 13.3 | 86.0 | 0.7 |
| % of all infections | 8.4 | 1.8 | 20.2 | 1.6 |
| Visits with RI, n | 6,437 | 1,221 | 5,161 | 55 |
| % of RI visits | 100.0 | 19.0 | 80.2 | 0.9 |
| % of all infections | 21.0 | 6.5 | 47.4 | 5.1 |

GII – gastrointestinal infections; RI – respiratory infections; Malaria – positivity by blood smear test with/without additional clinical symptoms, includes both severe (SM) and non-severe malaria (NSM).

The most likely co-infection combination was malaria and RI, with 5.3% of routine and 8.2% of walk-in visits, and the least likely co-infection combination was malaria and GII, with 0.8% of routine and 1.6% of walk-in visits (**Table S1**). Logistic regression analyses showed that infections tended to be non-redundant, and any infection significantly decreased the risk of any other infection reported at the same visit (**Table S2**).

### Distribution of the HBB, IFNL4 and IFNL3 polymorphisms in the Malian children and the populations of the 1000 Genomes Project

The *HbB*-A/A genotype, representing wild-type hemoglobin, was present in 80.5% of children; sickle cell trait (SCT, HbS-A/S or HbC-A/C) – in 18.6%, and sickle cell disease – (SCD, S/S, S/C or C/C) - in 0.9% of the children. The *IFNL4*-rs368234815 alleles (dG-72.2% and TT-28.8%) and *IFNL3*-rs4803217 alleles (G-66.7% and T-33.3%) were in moderate linkage disequilibrium (LD) in the total set of 914 Malian children (D’=0.98 and r^2^=0.78), which was also reflected in the haplotype frequencies of these variants (**Figure S3**). The allele and genotype frequencies of *HBB, IFNL3* and *IFNL4* polymorphisms were comparable to those in the populations of African ancestry in the 1000 Genomes Project, especially to the populations from West-Africa (**Table S3**). The distribution of alleles/genotypes of these variants in the four main tribes in Mali is also presented (**Table S3**).

### Associations between IFNL4 and IFNL3 polymorphisms and infection episodes in the Malian children

We analyzed the risk of an infection episode reported at any clinic visit from birth through up to 5 years of follow-up (ever/never analysis), and present the results adjusted for the tribes; unadjusted results are also presented in corresponding tables/figures. The rs368234815-dG allele was associated with an increased risk of GII (with a per-allele OR=1.53, 95%CI (1.13-2.07), p=0.005), malaria overall (OR=1.30, 95%CI (1.02-1.65), p=0.033), NSM (OR=1.28, 95%CI (1.00-1.63), p=0.047), but not of RI (OR=1.20, 95%CI (0.63-2.29, p=0.58) (Table 2).

**Table 2.**
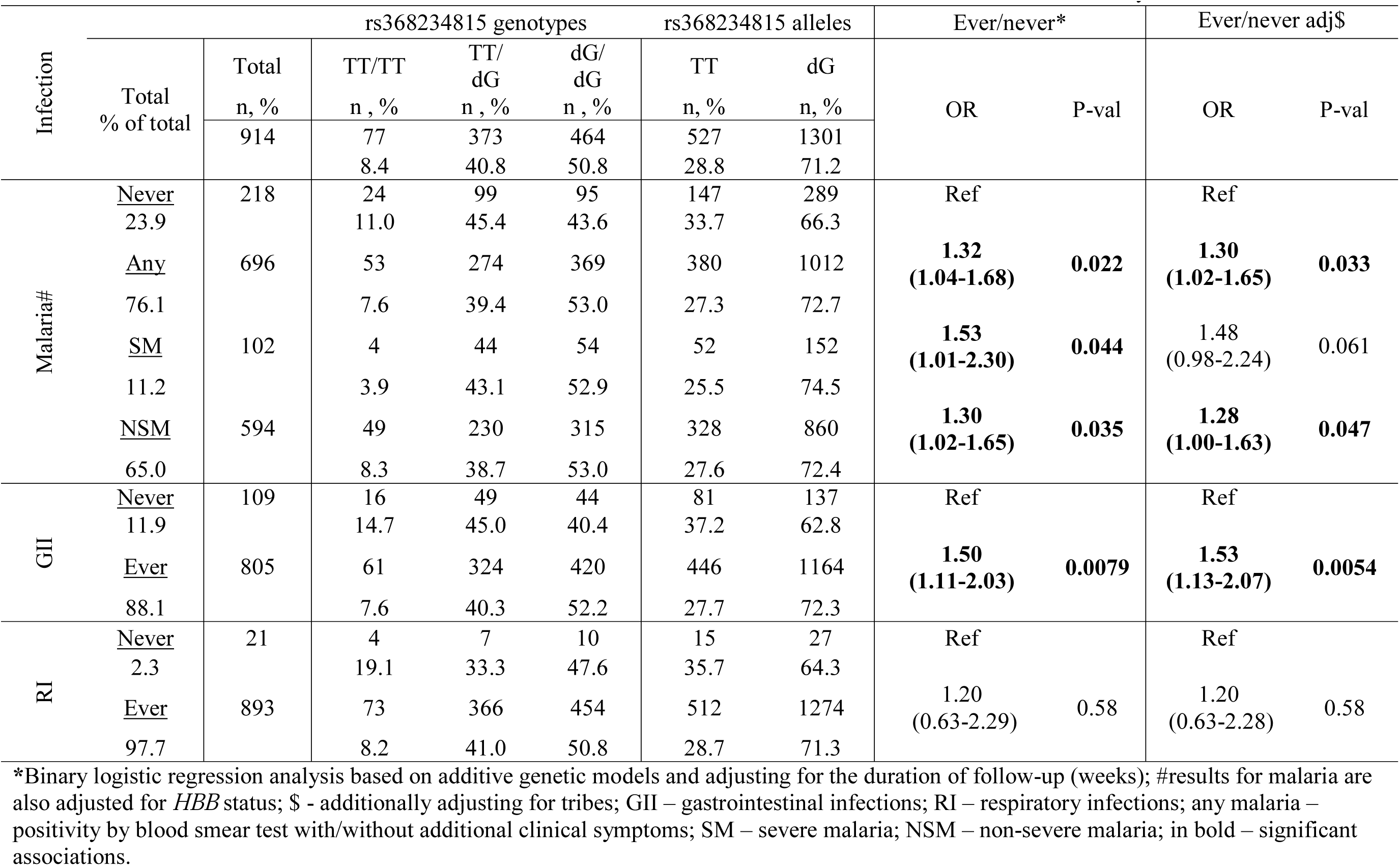
Associations between *IFNL4*-rs368234815 and infections in 914 children from the Mali birth cohort study

Gastrointestinal and respiratory symptoms might overlap with clinical manifestations of malaria. However, the associations of *IFNL4*-rs368234815 with these infections appeared independent, as mutual conditioning on these infections did not significantly alter the results (**Table S4**). We also analyzed the association between *IFNL4*-rs368234815 and time-to-first episode of each infection, which represents the earliest occurrence of symptoms of sufficient severity to require a visit to the clinic. The most significant association was observed for the *IFNL4*-rs368234815 and the time to the first episode of GII (per allele p=0.003) (Figure 1). The association was marginally significant for RI (p=0.045), but not for malaria overall and SM (Figure 1). The duration of follow-up did not differ significantly for children with different *IFNL4* genotypes (**Table S5**).

**Figure 1.**
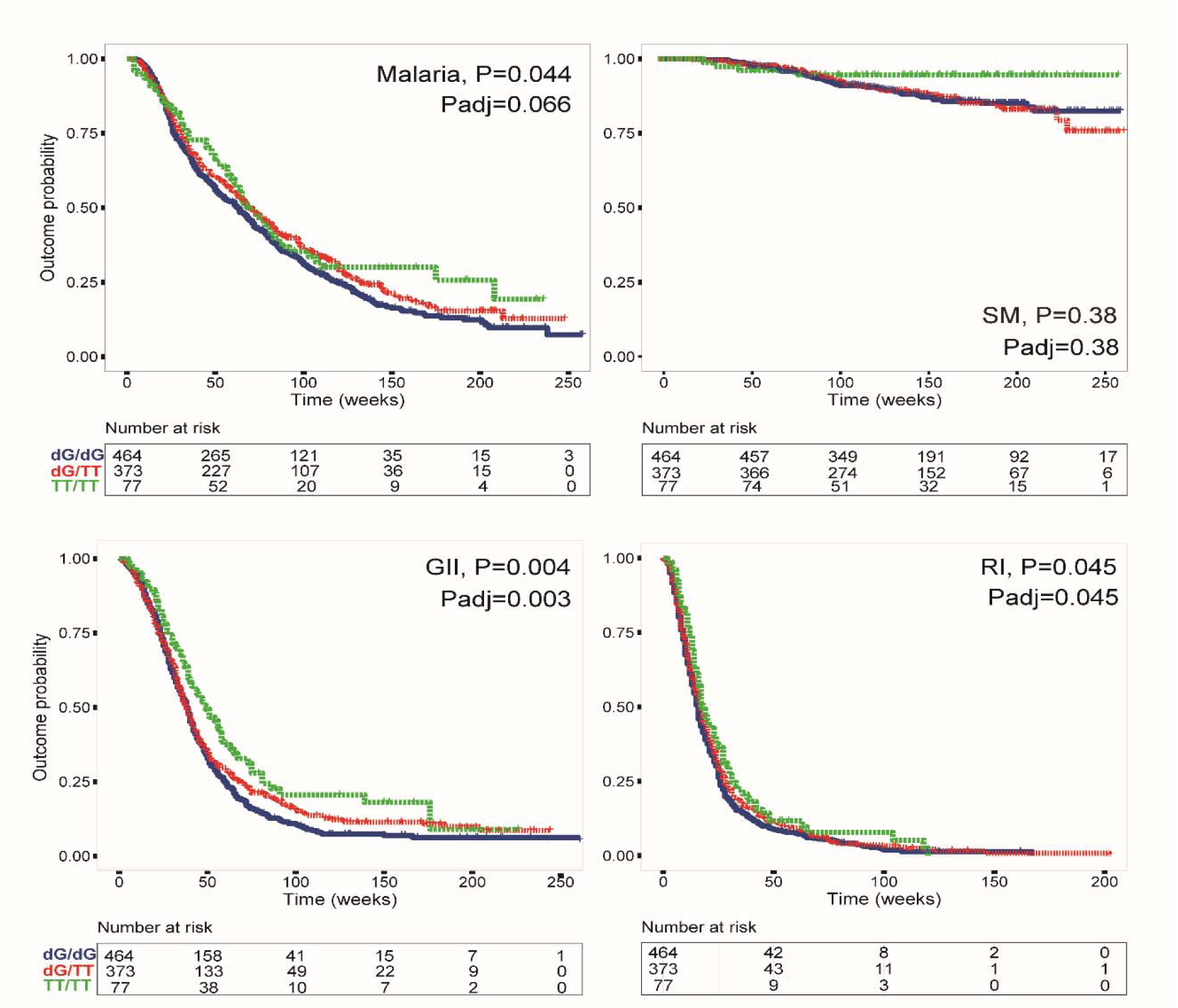
Time-to-first episode of infections in relation to *IFNL4*-rs368234815 polymorphism in 914 children from the Mali birth cohort study. Malaria – positivity by blood smear test with/without additional clinical symptoms, includes both severe (SM) and non-severe malaria; GII – gastrointestinal infections; RI – respiratory infections. The plots are for Kaplan-Meyer analysis, P-values are for Cox proportional hazards regression models with per-allele linear trends, for malaria – adjusting for *HBB* status. Padj-values are additionally adjusting for tribes.

The number of episodes reported for each infection during follow-up was the lowest for children with the IFN-λ4-Null genotype, but this was significant only in the dominant genetic model for GII (p=0.03, **Table S6**), suggesting that the increased susceptibility is related to infection overall and not its recurrence. All the associations for *IFNL3*-rs4803217 were weaker than for rs368234815 (**Table S7**, **Figure S4**) and lost significance in the models that included both variants, while most of the associations for rs368234815 remained significant (Table 3).

**Table 3.** Associations between *IFNL4*-rs368234815 and *IFNL3*-rs4803217 with infections

| Infection | | *IFNL4*-rs368234815 | | *IFNL3*-rs4803217 | | Models with both variants\$ | | | |
| --- | --- | --- | --- | --- | --- | --- | --- | --- | --- |
|  |  | rs368234815 |  | rs4803217 |  | rs368234815 |  | rs4803217 |  |
|  |  | OR, 95%CI | P-val | OR, 95%CI | P-val | OR, 95%CI | P-val | OR, 95%CI | P-val |
| Malaria # | Never | Ref |  | Ref |  | Ref |  | Ref |  |
|  | Any | **1.30**<br>**(1.02-1.65)** | **0.033** | 1.19<br>(0.94-1.50) | 0.15 | **1.73**<br>**(0.96-3.10)** | **0.068** | 0.74<br>(0.42-1.31) | 0.30 |
|  | SM | **1.48**<br>**(0.98-2.24)** | **0.061** | **1.52**<br>**(1.01-2.27)** | **0.044** | 1.04<br>(0.37-2.88) | 0.94 | 1.46<br>(0.54-3.98) | 0.45 |
|  | NSM | **1.28**<br>**(1.00-1.63)** | **0.047** | 1.15<br>(0.91-1.46) | 0.25 | **1.87**<br>**(1.03-3.38)** | **0.039** | 0.67<br>(0.37-1.19) | 0.17 |
| GII | Never | Ref |  |  |  |  |  |  |  |
|  | Ever | **1.53**<br>**(1.13-2.07)** | **0.0054** | **1.35**<br>**(1.00-1.82)** | **0.047** | **2.34**<br>**(1.06-5.19)** | **0.036** | 0.63<br>(0.29-1.39) | 0.26 |
| RI | Never | Ref |  |  |  |  |  |  |  |
|  | Ever | 1.2<br>(0.63-2.28) | 0.58 | 1.24<br>(0.66-2.33) | 0.51 | 0.94<br>(0.23-3.90) | 0.93 | 1.31<br>(0.32-5.31) | 0.71 |

\*Binary logistic regression analysis based on additive genetic models and adjusting for the duration of follow-up (weeks); in bold – significant associations; #results for malaria are adjusted for *HBB* status, \$ - results for models including both variants; any malaria – positivity by blood smear test with/without additional clinical symptoms; SM – severe malaria; NSM – non-severe malaria. \$-additionally adjusting for tribes.

## DISCUSSION

In 2015, 5.9 million children died globally before the age of five, with 30% of these deaths being caused by pneumonia, malaria and diarrhea ^15^. Thus, it is critical to identify and understand the factors affecting susceptibility and the host response to these infections. We identified the genetic ability to produce IFN-λ4, a type III IFN, in carriers of the *IFNL4*-rs368234815-dG allele, as a factor most significantly increasing the risk of gastrointestinal infections and earlier occurrence of these infections in Malian children under five. We also observed evidence for higher risk of malaria infection and earlier episodes of respiratory infections in carriers of the same allele.

Episodes of acute gastroenteritis (diarrhea and vomiting) are commonly caused by infections with rotavirus (RoV) or norovirus (NoV). Rotavirus is considered the most important global cause of severe diarrhea in children under five. In 2016 alone, despite more than 28,000 deaths considered to be averted by vaccination for rotavirus, still, an estimated 128,500 children under five died of this infection ^16^. In Rwanda, rotavirus was detected in rectal swabs of 36.9% of children under five seeking clinical help for acute gastroenteritis ^17^.

Norovirus is the second leading cause of acute gastroenteritis, accounting for up to 20% of all acute infections in all age groups worldwide ^18^. Although these infections are ubiquitous and in healthy individuals spontaneously clear within days, viral clearance may take longer in younger children, especially in those affected by malnutrition and other infections.

Extended severe dehydration due to diarrhea can be fatal, especially in infants. Human enteric viruses are difficult to study *in vitro*, but recent advances in human intestinal organoid models ^19,20^ should facilitate these studies. In contrast to humans that can produce either 3 or 4 type III IFNs (IFN-λ1, 2, 3 in all and IFN-λ4 - in a subset of individuals), the mouse genome invariably encodes only IFN-λ2 and IFN-λ3, which are collectively referred to as IFN-λ. Although most conclusions on RoV and NoV infections have been drawn using murine viruses and based on mouse models that differ from human cells by the IFN-λ repertoire, these studies proposed that intact signaling through type III IFNs is important for the control of GII. These conclusions were supported by the functional impact caused by the elimination of IFNLR1, which is a receptor expressed by epithelial cells and specific to the type III IFN family ^21–26^. However, the IFN-λ-related immune responses to enteric viruses appeared to be viral-strain and mouse-model specific ^22^, and could also be affected by the intestinal microbiota ^26,27^. Overall, clearance or persistence of enteric viruses likely results from the combination of factors contributed both by the host and the pathogen.

Although data on GII etiology was not available in our study, a large subset of these acute gastroenteritis episodes are likely to be caused by RoV and NoV - RNA viruses already linked with type III IFNs in animal models or *in vitro* studies. A higher risk and earlier episodes of clinical GII in children with *IFNL4*-rs368234815-dG allele might indicate an impaired immune response in these children due to the production of IFN-λ4. However, if diarrhea can be considered as a mechanism to purge those pathogens from the GI tract, these results might also indicate an enhanced innate immune response, as has been suggested based on mouse studies ^16^ and Kotenko, personal communication. The immune response altered in children with *IFNL4*-rs368234815-dG allele might manifest in more severe episodes od diarrhea but our study did not have information on the duration and the intensity of each infection episode. Either impaired or enhanced, the innate immune response associated with the genetic ability to produce IFN-λ4 appears to have negative health consequences. With the associations for GII being in the same direction as was previously reported for HCV and respiratory viruses, GII can be added to the list of outcomes negatively affected by the ability to produce IFN-λ4.

Acute RI cause 1.9 million deaths annually in children under five, with 70% of these deaths occurring in Africa and Southeast Asia ^28^. Several studies have explored the role of type III IFNs in the immune response to the respiratory syncytial virus (RSV) and influenza in the human and murine nasal epithelium. These studies suggested that type III IFNs represent the first line of defense from respiratory RNA viruses, and treatment with recombinant IFN-λs can protect from RI and stop the viral spread from upper to lower respiratory tract ^29–32^. Recently, a genetic association was reported for *IFNL4*-rs12979860-T and rs368234815-dG alleles with impaired clearance of respiratory RNA viruses in nasal swabs of children from Rwanda ^11^. The observed effect was in the same direction as for clearance of HCV reported in many studies, suggesting that the negative impact of IFN-λ4 is relevant for the clearance of respiratory RNA viruses as well. In our study, we observed a borderline significant association (p=0.045) with an earlier occurrence of RI in children with rs368234815-dG allele but the association with increased susceptibility to RI in these children did not reach statistical significance. This could be because respiratory symptoms recorded during clinic visits in our study were very common (were experienced by 97.7% of children during follow-up), and occurred very early in life, with the median timing of the first RI episode being at 16 weeks of age. Furthermore, RI episodes were defined very broadly, without stratifying by specific viral etiology, as was done in the study in Rwanda ^11^. In addition to bacterial and viral etiology, respiratory symptoms might also be caused by helminths (pinworms, roundworms, etc) transiting through the lungs during their life cycle. A more specific definition of RI etiology and surveillance of clearance is needed to clarify the association with genetic variants in the *IFNL3*/*IFNL4* region.

Our results showing the association between type III IFNs and malaria infection are also novel and, if validated, might present new mechanisms. Although III IFNs are mostly recognized for their antiviral role, the involvement of IFN-λs in bacterial and fungal infections have also been reported ^33^. Malaria is caused by plasmodium, an intracellular parasite, which first invades hepatocytes and then propagates in the liver where type III IFNs can be expressed after induction by different stimuli. However, these results were attenuated by adjusting for the tribe, suggesting possible differences between these tribes, such as malaria exposure and protection, that should be carefully considered as confounding factors in genetic association studies. Most of our results remained significant after adjustment for the tribe, warranting further replication studies.

While rs4803217 was previously suggested as a functional variant affecting the stability of *IFNL3* mRNA and possibly accounting for the association in the *IFNL3/IFNL4* region ^10^, this SNP appeared to be associated with all the outcomes in our study weaker than rs368234815, supporting *IFNL4*-rs368234815 as a primary associated and functional genetic variant in this region.

The strength of our study is the longitudinal analysis of a cohort of children who were regularly examined from birth to up to 5 years of age, a large sample size with 30,626 clinic visits that captured most, if not all, episodes of common infections. Longitudinal studies of transient but recurrent infectious diseases with high spontaneous clearance are more informative than cross-sectional studies, which may miss a significant proportion of events not detectable at the time of the study. The limitations of our study include a broad definition of gastrointestinal and respiratory symptoms without specifying their infectious etiology. Due to the regular medical assistance provided to the children at the time of the routine and walk-in visits, the negative health impacts of these infections were likely to be diminished, which makes our findings conservative. Even though the rs368234815-dG allele was associated with a higher risk of these infections in early childhood, our study did not explore the long-term health consequences of these associations. For example, we were not able to test whether earlier exposure to infections would affect immune fitness later in life. Additionally, our results do not explain why the ability to produce IFN-λ4 could be beneficial in Africa, accounting for its very high frequency, while this allele being selected against outside of Africa. Our exploratory study tested associations between multiple infections and models, which might result in some of these results being false-positive due to multiple testing and unadjusted confounding factors. Further studies in comparable cohorts are needed to confirm our findings and refine the etiology of these and other infections in different environmental and socio-economic settings.

In conclusion, our study suggests that type III IFNs, and IFN-λ4, specifically, negatively affect the immune response to infections in young children in Africa. Clearance of several RNA viruses likely impaired by IFN-λ4 (HCV, respiratory viruses, RoV and NoV) suggests that response to other clinically relevant RNA viruses, such a coronavirus, including SARS-CoV-2 that causes COVID-19, might also be impaired in carriers of the *IFNL4*-rs368234815-dG allele. These functional associations might contribute to the variable severity of clinical manifestations and outcomes in the infected patients, warranting further investigations.

## ACKNOWLEDGMENTS

We thank the study participants, the populations and health staff in the study area for their cooperation. We also thank the Mali Service Center for providing administrative support to the project, Drs. Richard Sakai and Souleymane Karambe for logistical support. We thank Dr. Sergei Kotenko, Rutgers University and Thomas O’Brien, Infections and Immunoepidemiology Branch, Division of Cancer Epidemiology and Genetics, National Cancer Institute for critical comments and discussions.

## Author contributions

LPO, SMM and PED conceived the idea and supervised the study; AM, OA, YS, JK, AD, RDM, MF and PED provided samples and data; AO, AAH and MM performed DNA extraction and genotyping of the samples; SK, PSA, RDM, OFV and LPO analyzed the data and generated figures and tables; LPO drafted the manuscript; RDM, OOO, NB, AD, SMM and PED helped with critical editing. All authors contributed to the interpretation of the results and read and approved the final manuscript.

