## Supplementary materials for "IFN-λ4 is associated with increased risk and earlier occurrence of gastrointestinal, respiratory and malarial infections in Malian children"

**Table S1**. Analysis of clinic visits reporting co-infections in 914 children from the Mali birth cohort study

| Visit types | **Visits, N** | **Malaria & GII** | **Malaria & RI** | **GII & RI** |
|---|---|---|---|---|
| Routine visits with at least 1 infection | 2377 | 19 (0.8%) | 127 (5.3%) | 58 (2.4%) |
| Walk-in visits with at least 1 infection | 8303 | 136 (1.6%) | 677 (8.2%) | 444 (5.3%) |

**Table S2.** Probability of co-infections reported at clinic visits, Log Odds in 914 children from the Mali birth cohort study

**Routine visits with at least one infection**

| | Second infection | | |
|---|---|---|---|
| First infection | Malaria | GII | RI |
| Malaria | | -2.79 | -3.35 |
| GII | -2.79 | | -1.88 |
| RI | -3.35 | -1.88 | |

**Walk-in visits with at least one infection**

| | Second infection | | |
|---|---|---|---|
| First infection | Malaria | GII | RI |
| Malaria | | -2.03 | -1.79 |
| GII | -2.03 | | -2.60 |
| RI | -1.79 | -2.6 | |

All logistic regression tests reported P-values < 2e-16 for Log Odds

**Table S3.** Distribution of *IFNL4*-rs368234815, *IFNL3*-rs4803217 and *HBB* polymorphisms in populations of African ancestry from the 1000 Genomes Project and 914 children from the Mali birth cohort study

| Gene | Alleles | ACB | ASW | LWK | ESN | GWD | MSL | YRI | All | Bamana | Peulh/ Fulani | Soninke | Malinke |
|---|---|---|---|---|---|---|---|---|---|---|---|---|---|
| Variant | Genotypes | Caribbean N=96 | USA N=61 | Kenya N=99 | Nigeria N=99 | The Gambia N=113 | Sierra Leone N=85 | Nigeria N=108 | Mali N=914 | Mali N=666 | Mali N=84 | Mali N=55 | Mali N=46 |
| ***IFNL4*** | TT | 0.26 | 0.3 | 0.44 | 0.28 | 0.22 | 0.29 | 0.27 | 0.29 | 0.28 | 0.36 | 0.27 | 0.29 |
| **rs11322783** | dG | 0.75 | 0.7 | 0.56 | 0.72 | 0.78 | 0.71 | 0.73 | 0.71 | 0.72 | 0.64 | 0.73 | 0.71 |
| corresponds to | TT/TT | 0.08 | 0.1 | 0.14 | 0.09 | 0.05 | 0.07 | 0.09 | 0.08 | 0.08 | 0.11 | 0.07 | 0.04 |
| rs368234815 | TT/dG | 0.34 | 0.41 | 0.61 | 0.38 | 0.34 | 0.44 | 0.35 | 0.41 | 0.39 | 0.50 | 0.39 | 0.50 |
| in 1000 Genomes | dG/dG | 0.57 | 0.49 | 0.25 | 0.53 | 0.61 | 0.49 | 0.56 | 0.51 | 0.53 | 0.39 | 0.54 | 0.46 |
| ***IFNL3*-** | G | 0.29 | 0.34 | 0.47 | 0.28 | 0.3 | 0.38 | 0.33 | 0.33 | 0.33 | 0.42 | 0.27 | 0.29 |
| **rs4803217** | T | 0.71 | 0.66 | 0.53 | 0.72 | 0.7 | 0.62 | 0.67 | 0.67 | 0.67 | 0.58 | 0.73 | 0.71 |
| | G/G | 0.1 | 0.15 | 0.18 | 0.07 | 0.09 | 0.15 | 0.12 | 0.11 | 0.11 | 0.15 | 0.07 | 0.04 |
| | T/G | 0.52 | 0.48 | 0.24 | 0.52 | 0.5 | 0.39 | 0.45 | 0.45 | 0.44 | 0.54 | 0.40 | 0.50 |
| | T/T | 0.38 | 0.37 | 0.58 | 0.41 | 0.41 | 0.46 | 0.43 | 0.44 | 0.45 | 0.31 | 0.53 | 0.46 |
| ***LD* between markers** | D’ | 0.97 | 1.00 | 0.85 | 0.92 | 1.00 | 1.00 | 1.00 | 0.98 | 0.98 | 1.00 | 0.95 | 1.00 |
| | $r^2$ | 0.78 | 0.86 | 0.66 | 0.83 | 0.67 | 0.65 | 0.73 | 0.78 | 0.76 | 0.76 | 0.91 | 1.00 |
| ***HBB*** | T | 0.95 | 0.98 | 0.9 | 0.88 | 0.89 | 0.88 | 0.86 | 0.996 | 0.996 | 0.994 | 0.991 | 0.989 |
| **rs334** | A | 0.05 | 0.02 | 0.1 | 0.12 | 0.12 | 0.12 | 0.14 | 0.004 | 0.004 | 0.006 | 0.009 | 0.011 |
| | T/T | 0.91 | 0.97 | 0.8 | 0.76 | 0.77 | 0.75 | 0.72 | 0.91 | 0.90 | 0.95 | 0.91 | 0.87 |
| | A/T | 0.09 | 0.03 | 0.2 | 0.24 | 0.23 | 0.25 | 0.28 | 0.09 | 0.096 | 0.048 | 0.091 | 0.13 |
| | A/A | 0 | 0 | 0 | 0 | 0 | 0 | 0 | 0.0011 | 0.002 | 0 | 0 | 0 |
| ***HBB*** | G | 0.96 | 0.98 | 1.0 | 1.0 | 1.0 | 0.99 | 0.97 | 0.95 | 0.948 | 0.976 | 0.955 | 0.967 |
| **rs33930165** | A | 0.04 | 0.02 | 0 | 0 | 0.004 | 0.006 | 0.030 | 0.050 | 0.052 | 0.024 | 0.045 | 0.033 |
| | G/G | 0.93 | 0.97 | 1.0 | 1.0 | 0.99 | 0.99 | 0.94 | 0.90 | 0.90 | 0.95 | 0.93 | 0.94 |
| | G/A | 0.07 | 0.03 | 0 | 0 | 0.009 | 0.010 | 0.060 | 0.096 | 0.104 | 0.048 | 0.055 | 0.065 |
| | A/A | 0 | 0 | 0 | 0 | 0 | 0 | 0 | 0.001 | 0 | 0 | 0.018 | 0 |

*HBB* status: WT – wild-type; SCT – sickle cell trait; SCD – sickle cell disease; 1000 Genomes Project populations of African ancestry (AFR) include Yoruba in Ibadan Nigeria (YRI), Luhya in Webuye, Kenya (LWK), Mandinka in The Gambia (MAG), Mende in Sierra Leone (MSL), Esan in Nigeria (ESN), Americans of African Ancestry in South-West USA (ASW), and African Caribbean in Barbados (ACB), based on information from https://www.internationalgenome.org/1000-genomes-browsers/. LD between markers was analyzed with LDLink (https://ldlink.nci.nih.gov/?tab=home). Mali dataset includes data for all 914 children and the four major tribes.

**Table S4.** Associations between *IFNL4*-rs368234815 and mutually adjusted risk of infections in 914 children from the Mali birth cohort study

| Infection | Total<br>% of total | Ever/never* | | Ever/never, adjusting for episodes of other infections | | | | | |
|---|---|---|---|---|---|---|---|---|---|
| | | OR (95%CI) | P-val | OR (95%CI) | P-val | OR (95%CI) | | OR (95%CI) | P-val |
| Malaria# | Never<br>23.9 | Ref | | Ref | | Ref | | Ref | |
| | Any<br>76.1 | **1.32 (1.04-1.68)** | **0.022** | **1.31 (1.03-1.66)** | **0.030** | **1.3 (1.02-1.65)** | **0.034** | **1.31 (1.03-1.66)** | **0.031** |
| | | | | Adjusted for GII | | Adjusted for RI | | Adjusted for GII and RI | |
| GII | Never<br>11.9 | Ref | | Ref | | Ref | | Ref | |
| | Ever<br>88.1 | **1.5 (1.11-2.03)** | **0.0079** | **1.54 (1.14-2.09)** | **0.0050** | **1.49 1.1-2.01** | **0.0076** | **1.50 1.11-2.03** | **0.0069** |
| | | | | Adjusted for malaria | | Adjusted for RI | | Adjusted for malaria and RI | |
| RI | Never<br>2.3 | Ref | | Ref | | Ref | | Ref | |
| | Ever<br>97.7 | **1.2 0.63-2.29** | 0.58 | 1.2 (0.63-2.29) | 0.58 | 1.12 0.59-2.13 | 0.74 | 1.12 0.59-2.14 | 0.73 |
| | | | | Adjusted for malaria | | Adjusted for GII | | Adjusted for malaria and GII | |

All the results are also adjusted for the tribes.

**Table S5.** Time of follow-up (weeks) from birth in relation to *IFNL4*-rs368234815 in 914 children from the Mali birth cohort study

| All children, weeks of follow-up Median (95% CI) | *IFNL4*-rs368234815 genotypes, weeks of follow-up, Median (95% CI) | | | P-value for non-parametric one-way ANOVA, Kruskal-Wallis test |
|---|---|---|---|---|
| | dG/dG | dG/TT | TT/TT | |
| 140.0 (143.5-150.5) | 142.5 (144.0-154.0) | 140 (140.4-151.2) | 143 (127.8-154.3) | 0.537 |
| Min-max time range, weeks 52-264 | 52-262 | 52-264 | 53-261 | |

**Table S6**. Relative rates of infection episodes per year in relation to *IFNL4*-rs368234815 genotypes during 1-5 years of follow-up in 914 children from the Mali birth cohort study

| Infection | Max episode counts, N | Geometric mean*, N | | | | Additive model#, Per-dG allele | | Dominant model# dG/dG and dG/TT vs. TT/TT | |
|---|---|---|---|---|---|---|---|---|---|
| | | All genotypes | dG/dG | dG/TT | TT/TT Ref | Relative rate | P-value | Relative rate | P-value |
| Malaria | 22 | 2.82 | 2.84 | 2.86 | 2.45 | 1.08 | 0.14 | 1.17 | 0.21 |
| SM | 3 | 1.18 | 1.23 | 1.13 | 1.19 | 1.22 | 0.21 | 2.13 | 0.14 |
| NSM | 22 | 2.75 | 2.76 | 2.79 | 2.44 | 1.07 | 0.18 | 1.15 | 0.28 |
| GII | 14 | 2.49 | 2.51 | 2.54 | 2.20 | 1.05 | 0.23 | 1.22 | **0.03** |
| RI | 25 | 5.33 | 5.43 | 5.30 | 4.80 | 1.01 | 0.79 | 1.05 | 0.47 |

Malaria – includes both severe (SM) and non-severe malaria (NSM); GII – gastrointestinal infections; RI – respiratory infections. For malaria episodes – adjusted for *HBB* status. Geometric means were calculated excluding children who never experienced an infection episode (zero episodes). *Geometric mean is computed by exponentiating the average log-transformed number of episodes. The relative rate represents an increase in the number of episodes per year compared to the reference genotype group (TT/TT, IFN-λ4-Null), according to additive or dominant genetic models, and adjusting for the tribes.

**Table S7.** Associations between *IFNL3*-rs4803217 and infections in 914 children from the Mali birth cohort study

| | | | rs4803217 genotypes | | | rs4803217 alleles | | Ever/never* | | Ever/never$ | |
|---|---|---|---|---|---|---|---|---|---|---|---|
| Infections | Total | Total | G/G | T/G | T/T | OR | OR | OR | P-val | OR | P-val |
| | % of total | n, % | n , % | n , % | n , % | n, % | n, % | | | | |
| | | 914 | 101 | 407 | 406 | 609 | 1219 | | | | |
| | | | 11.5 | 44.5 | 44.4 | 33.3 | 66.7 | | | | |
| Malaria# | Never | 218 | 29 | 102 | 87 | 160 | 276 | Ref | | Ref | |
| | 23.9 | | 13.3 | 46.8 | 39.9 | 36.7 | 63.3 | | | | |
| | Any | 696 | 72 | 305 | 319 | 449 | 943 | 1.2 | 0.122 | 1.19 | 0.15 |
| | 76.1 | | 10.3 | 43.8 | 45.8 | 32.3 | 67.7 | (0.95-1.52) | | (0.94-1.50) | |
| | SM | 102 | 4 | 48 | 50 | 56 | 148 | **1.56** | **0.032** | **1.52** | **0.044** |
| | 11.2 | | 3.9 | 47.1 | 49 | 27.5 | 72.5 | **(1.04-2.33)** | | **(1.01-2.27)** | |
| | NSM | 594 | 68 | 257 | 269 | 393 | 795 | 1.16 (0.92-1.47) | 0.22 | 1.15 (0.91-1.46) | 0.25 |
| | 65 | | 11.4 | 43.3 | 45.3 | 33.1 | 66.9 | | | | |
| GII | Never | 109 | 16 | 54 | 39 | 86 | 132 | Ref | | | |
| | 11.9 | | 14.7 | 49.5 | 35.8 | 39.4 | 60.6 | | | | |
| | Ever | 805 | 85 | 353 | 367 | 523 | 1087 | 1.34 (0.99-1.80) | 0.055 | **1.35 (1.00-1.82)** | **0.047** |
| | 88.1 | | 10.6 | 43.9 | 45.6 | 32.5 | 67.5 | | | | |
| RI | Never | 21 | 4 | 9 | 8 | 17 | 25 | Ref | | | |
| | 2.3 | | 19 | 42.9 | 38.1 | 40.5 | 59.5 | | | | |
| | Ever | 893 | 97 | 398 | 398 | 592 | 1194 | 1.24 | 0.51 | 1.24 | 0.51 |
| | 97.7 | | 10.9 | 44.6 | 44.6 | 33.1 | 66.9 | (0.66-2.33) | | (0.66-2.33) | |

*****Binary logistic regression analysis based on additive genetic models and adjusting for the duration of follow-up (weeks); #results for malaria are also adjusted for *HBB* status; $ - additionally adjusting for the tribes; GII – gastrointestinal infections; RI – respiratory infections; any malaria – positivity by blood smear test with/without additional clinical symptoms; SM – severe malaria; NSM – non-severe malaria; in bold – significant associations.

### SUPPLEMENTARY FIGURES

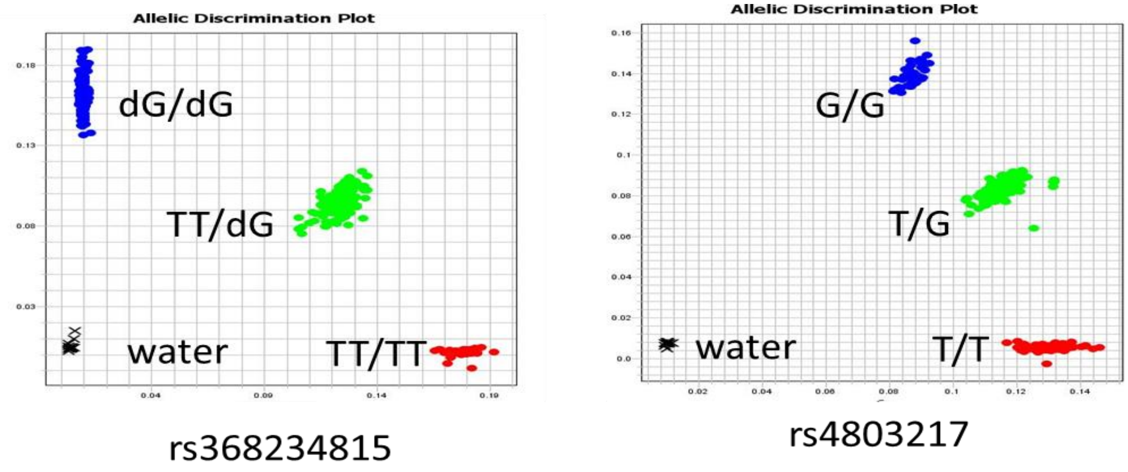


**Figure S1**. Representative allelic discrimination plots for genotyping of *IFNL4*-rs368234815 and *IFNL3*-rs4803217 polymorphisms in the Mali birth cohort study samples by custom TaqMan genotyping assays.

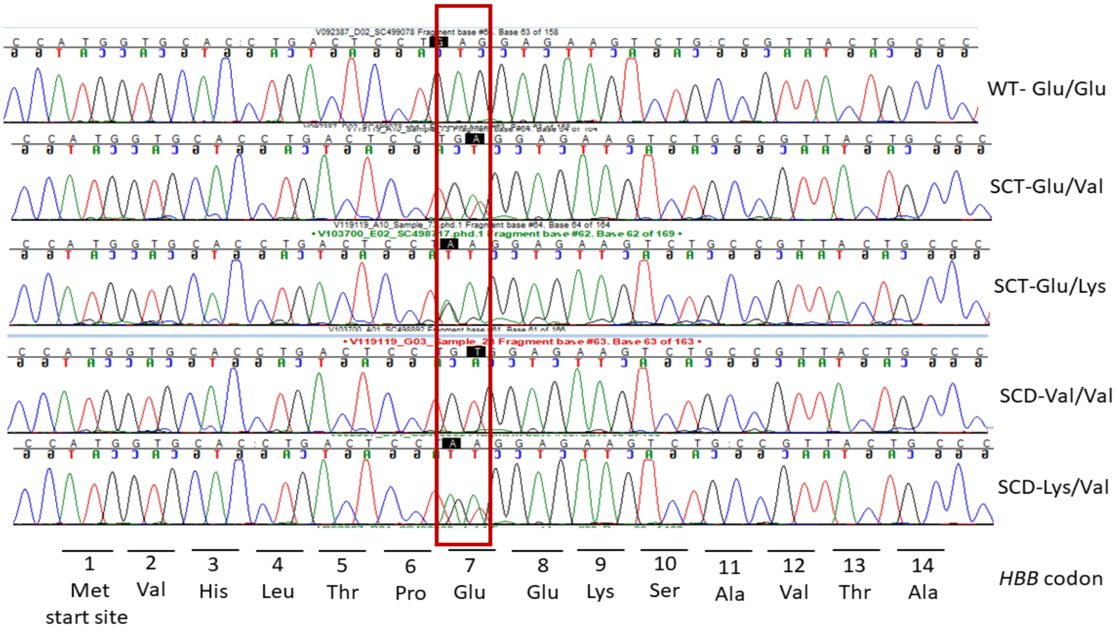


**Figure S2.** Representative Sanger sequencing chromatograms for genotyping of the *HBB* polymorphisms rs33930165-A/G (Glu7Lys) and rs334-A/T (Glu7Val) in the Mali birth cohort study samples, as has been previously described (13).

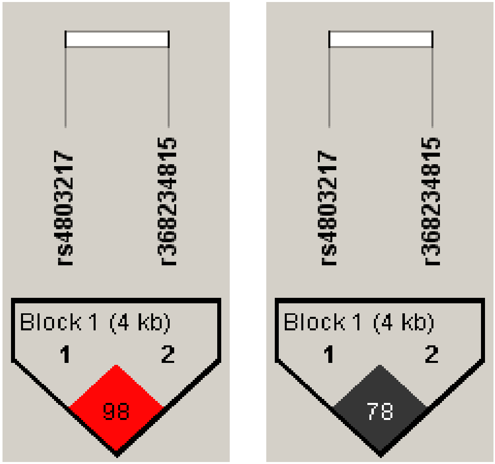


D’ r2

| **Haplotypes of *IFNL4*-rs368234815 and *IFNL3*-rs4803217 in 914 children from the Mali birth cohort study** | | | | |
|---|---|---|---|---|
| Haplotype | *IFNL4* frame-shift rs368234815 | *IFNL3* 3’UTR rs4803217 | Frequency, % | IFN-λ4 protein |
| 1 | dG | T | 66.4 | produced |
| 2 | TT | G | 28.5 | not produced |
| 3 | dG | G | 4.8 | produced |

**Figure S3**. HaploView linkage disequilibrium (LD) plots and haplotypes of the *IFNL4*-rs368234815 and *IFNL3*-rs4803217 polymorphisms in 914 children from the Mali birth cohort study. The plots show LD metrics - D’=0.98; $r^2$=0.78.

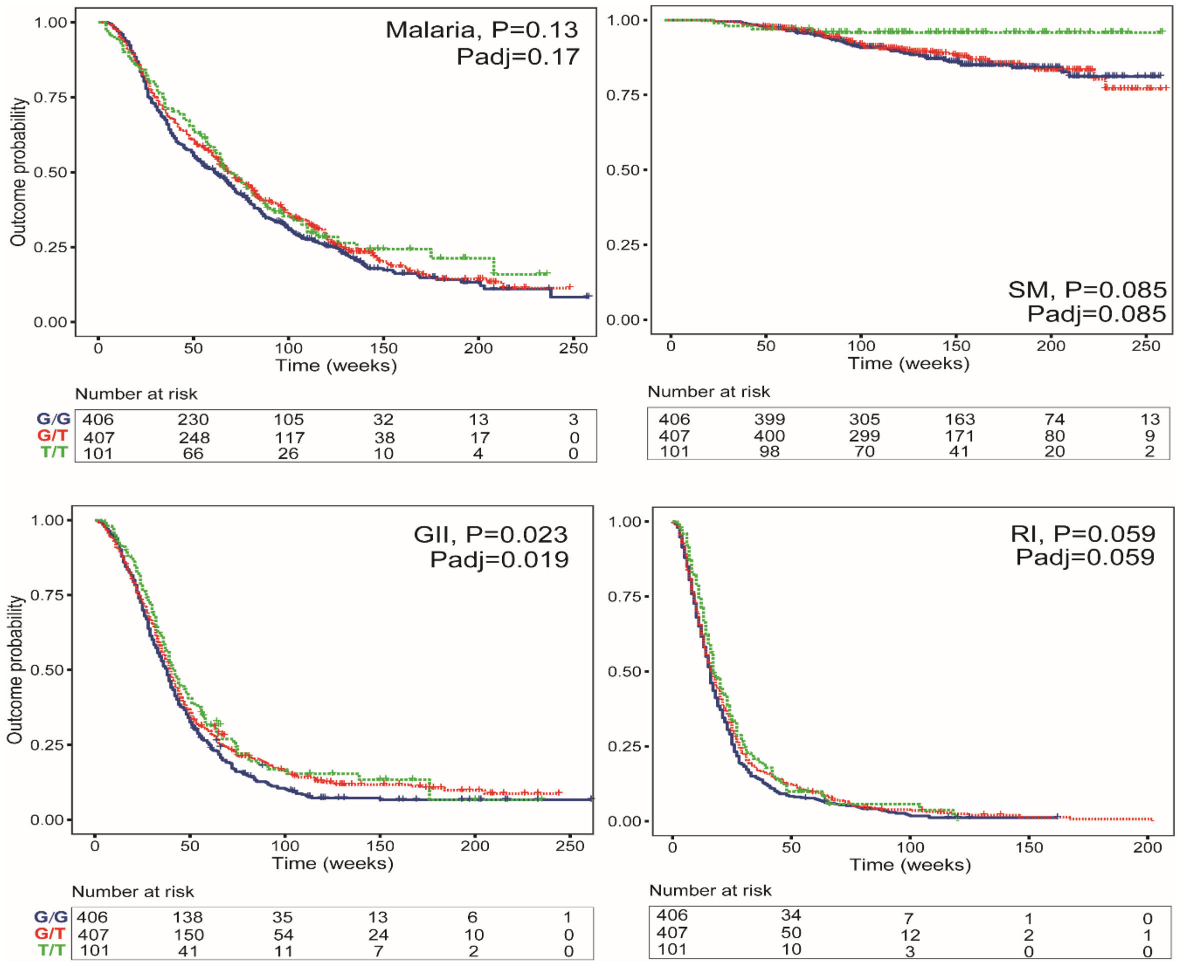


**Figure S4.** Time to the first episode of infections in relation to *IFNL3*-rs4803217 polymorphism in 914 children from the Mali birth cohort study. Malaria – positivity by blood smear test with/without additional clinical symptoms, includes both severe (SM) and non-severe malaria; GII – gastrointestinal infections; RI – respiratory infections. The plots are for Kaplan-Meyer analysis, P-values are for Cox proportional hazards regression models with per-allele linear trends, for malaria – adjusting for *HBB* status. Padj - additionally adjusting for the tribes.
